# Tree species identity drives soil Carbon and Nitrogen stocks in nutrient-poor sites

**DOI:** 10.1101/2023.05.15.540797

**Authors:** Estela Covre Foltran, Norbert Lamersdorf

## Abstract

The establishment of mixed forest stands can be seen as an option to enhance soil organic carbon stocks and to protect forest ecosystems from various impacts of climate change. Increasing temperatures and drought potentially affect the vitality of the native coniferous Norway spruce *(Picea abies)*, often used in mixed forests. We investigated the effects of a replacement of Norway spruce by Douglas fir (*Pseudotsuga menziesii*) admixed to European beech (*Fagus sylvatica*) on C and nitrogen (N) concentrations and stocks, as well as the vertical distribution and changes in forest floor and mineral soil (down to 30 cm depth). Each site included a quintet of neighboring forest stands of European beech, Douglas fir, and Norway spruce stands as well as mixtures of beech with either Douglas fir or spruce. The stands were located in two regions with different soil conditions (loamy vs sandy soils). Our results showed that the C stocks of the organic layer were significantly influenced by tree species, while the C stock of the mineral soil varied among soil types. Total soil organic C stocks demonstrated notable species-specific characteristics, primarily driven by the elevated C stocks in the organic layer. In sandy soils, conifers and mixed forests allocated 10% more C and N in the organic layer compared to loamy soils, whereas the C and N stocks under beech remained consistent, regardless of the site condition. The interaction between species and sites was significant only for Douglas fir and mixed Douglas fir/beech, indicating that the impact of species on C and N varied across sites and was notably pronounced in sandy soils. The higher potential for carbon and N storage in mixed-species forests compared to pure stands emphasizes the capacity of mixed forests to provide valuable ecosystem services, enhancing C sequestration in sandy soils.

## 1. Introduction

Forest soils are complex ecosystems and their proper management, especially through the selection of trees with distinct species identities, may have a major impact on soil organic carbon (SOC) stocks (Angst et al., 2018; Jandl et al., 2007; Vesterdal et al., 2013) leading to C sequestration potentials.

Typical driving factors affecting C and N storages by tree species identities include the leaf morphology, the quality of leaf litter, and the distinct rooting characteristics exhibited by different tree species (Bolte and Villanueva, 2006). Hence, the identity of tree species can significantly influence carbon (C) stocks and C:N ratios at the ecosystem level, especially within the organic layer and upper mineral soil layers (Dawud et al., 2017; Vesterdal et al., 2013). Furthermore, the species diversity drives the fluxes of carbon and nitrogen in soils and their role in C sequestration has been highlighted to improve the ecosystem functions (Cepáková et al., 2016; Mueller et al., 2012; Vesterdal et al., 2002). Different patterns of C storage in soil profile and organic layers have been reported for conifers and deciduous trees for temperate ecosystems (Bolte and Villanueva, 2006). The differences in litter input, root distribution (Oulehle et al., 2007) as well as decomposition rates (Cools et al., 2014) have been suggested as the most likely explanations for tree species influence on soil C stocks. In Central Europe, enrichment of European beech *(Fagus sylvatica)* stands with conifers results in mixtures that provide, in terms of wood production, sufficient economic returns (Kölling et al., 2009), and considering a structurally diverse forests represent an important element of approaches to deliver a wide range of ecosystem goods and services like carbon sequestration (Ammer, 2019). The conventional conifers managed are native Norway spruce *(Picea abies)* and the coastal provenance of the non-native Douglas fir *(Pseudotsuga menziesii var. menziesii)* (Neuner et al. 2015). However, Norway spruce in Europe is frequently affected by wind throw, bark beetle infestations, and climate change induced drought (Dobor et al., 2020; Hlásny and Turčáni, 2013; Kölling and Zimmermann, 2007). These susceptibilities raise questions concerning the enrichment of European beech stands with Norway spruce that have been promoted over past decades. Enrichment with coastal Douglas fir may be a more suitable option but not much is known on the effect of Douglas fir on ecosystem functioning (Glatthorn, 2021).

Often, Douglas fir and European beech show high fine root density in deeper soil layers (Bolte and Villanueva, 2006) increasing C allocation in mineral soil layers. In contrast, the native conifer Norway spruce (*Picea abies*), known as a shallow-rooted tree species, accumulates C preferentially in the upper mineral soil layers. Moreover, it has already been shown, that the admixtures of conifers (either with Douglas fir or Norway spruce) to beech forests modified the soil C and N input through the fine root distribution in the soil profile (Vesterdal and Raulund-Rasmussen, 1998). Additionally, the enhanced nutrient concentrations of beech litter increased litter decomposition of conifers needles (Krishna and Mohan, 2017; Neumann and Martinoia, 2002; Vesterdal et al., 2008), followed by less C accumulation in the organic soil layers and shift to potential stable C pools in the mineral soil. However, there is still inconsistent information on how mixed stands of beech with Douglas fir would affect the vertical distribution of forest soil carbon and nitrogen. Due to differences in litter input and decomposition rates, it is most likely that mixed stands enhance the organic carbon stocks in the organic layers. Thus, a deeper root system reported for Douglas fir and beech, along with a high C concentration in the organic layer, might suggest that the admixture of Douglas fir into beech stands potentially results in a high total carbon stocks throughout the entire soil profile.

Therefore, the main objective of our study was to analyze the accumulation, vertical distribution as well as litter quality indices (C:N ratio) of the upper mineral C and N stocks of different pure and mixed stand types (pure European beech, pure Norway spruce, pure Douglas-fir, mixed European beech/Norway spruce and mixed European beech/Douglas-fir) along northern Germany under distinct soil characteristics (loamy *versus* sandy soil conditions). We hypothesized i) the admixture of Douglas fir to beech forests will increase C stocks at the mineral soil compared to respectively beech monocultures, ii) Carbon stocks in the litter layer will diminish under the introduction of Douglas fir to beech forests, with a consequent transfer of carbon from the forest floor to more stable carbon pools such as the humus layer and upper mineral soil, compared to the respective monocultures, and iii) effects of species identity on C stocks will be more pronounced on nutrient-poor sites (sandy soils) than on nutrient-rich soils (loamy soil).

## 2. Material and Methods

### 2.1. Study sites

We investigated eight sites in Lower Saxony, northern Germany. Each site includes a quintet of neighboring forest stands. Three of these stands were monospecific stands of European beech (Be - *Fagus sylvatica*), Norway spruce (S - *Picea abies*), or Douglas fir (D - *Pseudotsuga menziesii*). The other two were mixed stands, one composed of spruce/beech (SB) and one composed of Douglas fir/beech (DB). Essential criteria for the selection of the sites were the presence of the respective quintets in a similar advanced silvicultural development status, altogether, we employed a multi-criteria approach, considering factors such as climate, soil type (based on German soil inventory), stand age, and existing research on land-use, as well as a relevant single tree admixture in the mixed stands.

However, on the basis of the available forest site mapping and also confirmed by Foltran et al. (2023), two groups with relatively homogeneous site conditions were distinguished in our data set, in particular via the texture data (Table 1; Fig. S3), but also addressing the geological substrate classes, the correspondently developed soil types as well as the given range of water and nutrient levels.

**Table1.**
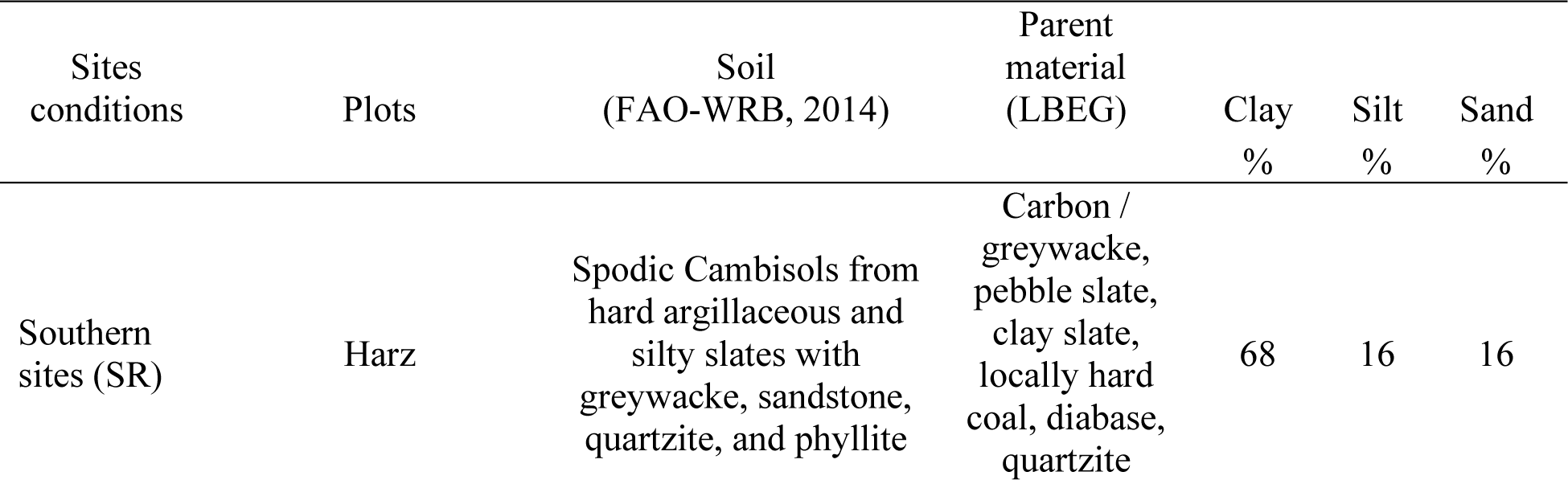
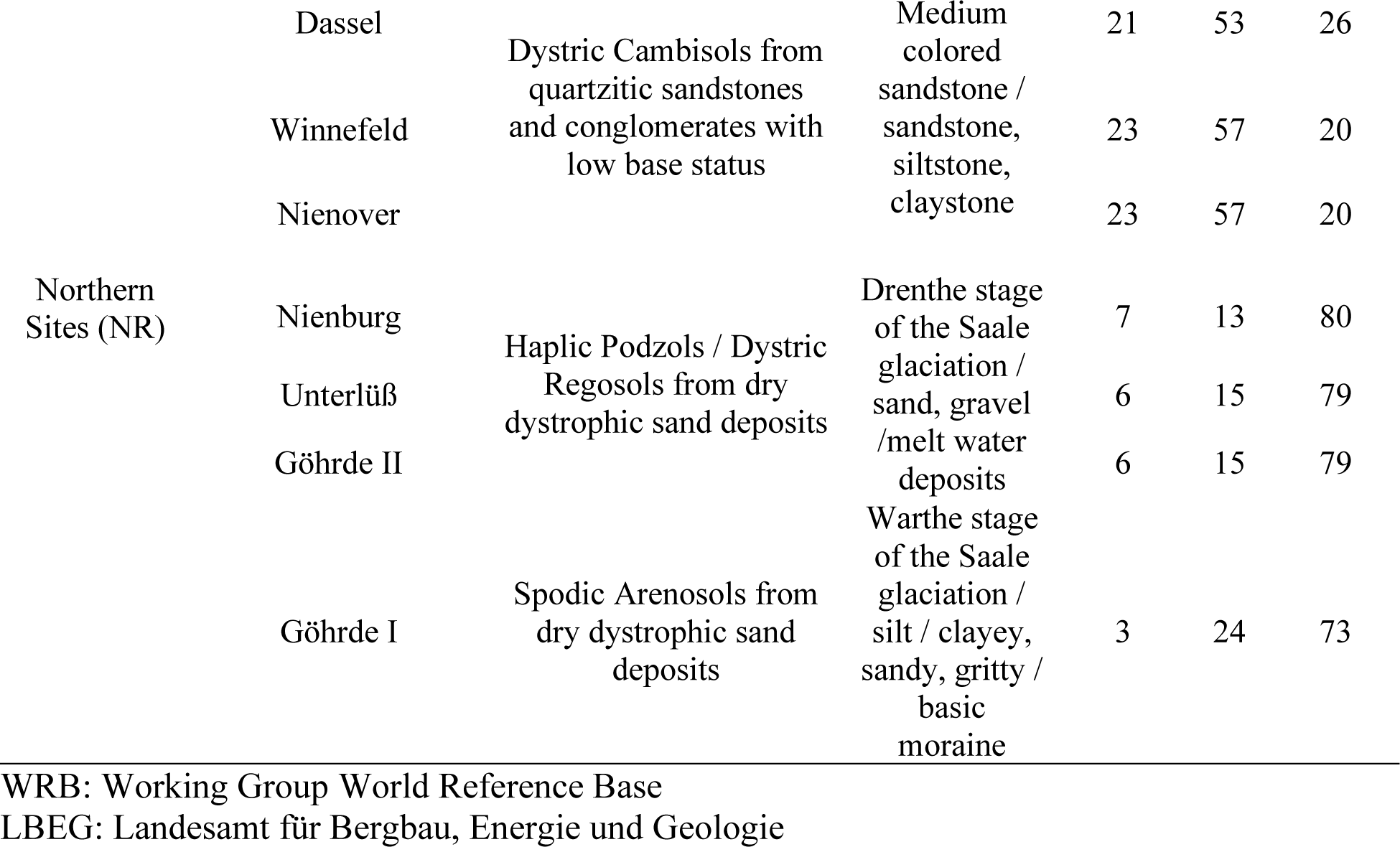
Soil classification from each plot is given following (FAO, 2014). Soil parent material was identified matching the plot coordinates with the German National inventory database (LBEG). Soil texture was measured by the integral suspension pressure method (ISP) and determined by PARIO.

The first group comprehended four sites in the South of Lower Saxony (Southern sites). The parental rock material is either loess-influenced Triassic sandstone or Paleozoic shares of greywacke, sandstone, quartzite, and phyllite, leading to soil types of partly Spodic Cambisols and Dystric Cambisols (FAO, 2014). The mean annual precipitation is 821 – 1029 mm (calculated from German Weather Service since 1981). The second group is formed by four sites at the North of Lower Saxony (Northern sites), located on out-washed sand with the Haplic Podzol and Spodic Arenosols soils (Table 1). The mean annual precipitation is 672 – 746 mm. Concerning nutrients and according to the official Lower Saxony forest site mapping and Foltran et al. (2023), the southern sites (loamy soils) are nutrient-rich, and the northern sites (sandy soils) are nutrient-poor.

### 2.2. Soil sampling

In each forest stand type (50 m x 50 m) at all sites 4 randomly selected points were chosen as representative sampling points. These selected points were oriented at the stand-level, e.g., we standardized two meters of minimal distance from the trees to avoid coarse roots. At each sampling plot, the forest floor was collected using a steel frame (*d*=28 cm) and sorted by identifiable foliar (Ol – Litter), non-foliar (Of – decay layer), and non-identifiable and humified (Oh – Humus) of the organic layer. Mineral soil was sampled using a core auger (*d*=8 cm), separated at 0-5, 5-10, and 10-30 cm soil depth. Bulk soil density from each depth was calculated using soil metal rings (250 cm³) to further stock analysis.

Partly missing bulk density data (n=140; 33% of total) due to frozen soil conditions, interfering tree roots or stones during sampling were estimated by Adamś equation (Adams, 1973) adapted by (Chen et al., 2017). The approach uses soil organic matter (SOM) and pH as bulk density predictors.

### 2.3. Sample preparation and analysis

All mineral soil samples were oven-dried at 40°C until constant weight and sieved through 2 mm mesh, subsamples from the fine soil fractions (<2 mm diameter) were ground with a Retsch mortar grinder RM 200 (Retsch, Germany) for 10 min. The organic layer samples were dried at 60°C until constant weight, weighted and ball milled (MM2, Fa Retsch) for further analysis.

All the samples were analyzed for total C and N by dry combustion in a Leco CSN 2000 analyzer. There was no inorganic C (CaCO_3_) within 30 cm depth in soils and all measured C were consequently considered to be organic.

The P from organic layers was determined by pressure digestion with 65% nitric acid for 8 h at 170°C (Höhle et al., 2018). The digestates were filtered by ash-free cellulose filters and determined by ICP-OES (Spectro Genesis).

The pH measurements for all mineral soil samples were performed in KCl solution, more detailed descriptions can be found in Foltran et al. (2023). Further details on the analytical procedure can be found in (König et al., 2014).

### 2.4. Calculations of Stocks

We estimated the soil bulk density from the oven-dried and moisture corrected (105 ◦C) fine soil mass and its volume. The fine soil volume was estimated from the difference between the volume of the soil corer and the volume of stones and roots. Carbon and N stocks in each layer were estimated from the organic layer dry weight (O-layers) and soil bulk density (mineral soil), concentrations of C and N and depth of the individual soil depths for each sampling point.

### 2.5. Statistical Analyses

To consider the potentially confounding effects of the parent material on topsoil conditions, we clustered our sites in two “soil type” groups based on soil texture (% Clay, % Silt, % Sand – Table 1). Moreover, we performed a principal component analysis (PCA) considering all available soil variables and sites (soil data from Foltran et al., 2023). In each biplot, 95% confidence level ellipses were added, grouping all sites (1-8), therefore, clustered into 2 distinctive groups, nutrient-rich sites (loamy soils) and nutrient-poor sites (sandy soils) (Fig. S1).

To address the non-independent nature of multiple horizons within one soil profile for this subset of data, we chose linear mixed effect models (Rasmussen et al., 2018). To estimate the effect of forest stand and site condition (loamy vs sandy), we fitted linear mixed models (LMMs) to log-transformed response variables (C and N) and then applied planned contrasts (Piovia-Scott et al., 2019). All LMMs included forest stands (European beech, Douglas fir, Douglas fir/beech, Norway spruce, and spruce/ beech), site conditions (loamy and sandy sites), and soil depths (Ol, Of, Oh [organic layers], and 0–5, 5–10, and 10–30 cm [mineral soil]) as fixed effects. The eight sites were included as random factor. Models were stepwise selected by likelihood ratio test, and minimal models included all main effects and the interaction of forest type and region. The model performances are available on the supplementary material.

Moreover, we used the ggscatterstats package (Patil, 2021) to visualize the relationship between *SOM x pH* and *N x pH*, using a linear regression with confidence interval (<0.95). All analyses were done in R 4.2.3. We used the ‘nlme’ package to fit LMMs and the ‘emmeans’ package for planned contrasts. All mixed models met the assumptions of normality of residuals and homogeneity of variance.

## 3. Results

### 3.1. Organic layer dry mass and soil density

No effects of the factor Site conditions on the organic layers dry masses were observed. Instead, the terms Depth (Dth), Forest stand (FS) and their respective interaction were consistency significant affecting the organic layer dry mass (Table 2 – Fig. S2), where both conifers (D and S) and the mixed stands (DB and SB) showed higher dry mass at the Oh layer than pure Be, meanwhile the Ol layer showed the lowest dry mass in mixed stands, independent of the site conditions.

**Table 2.**
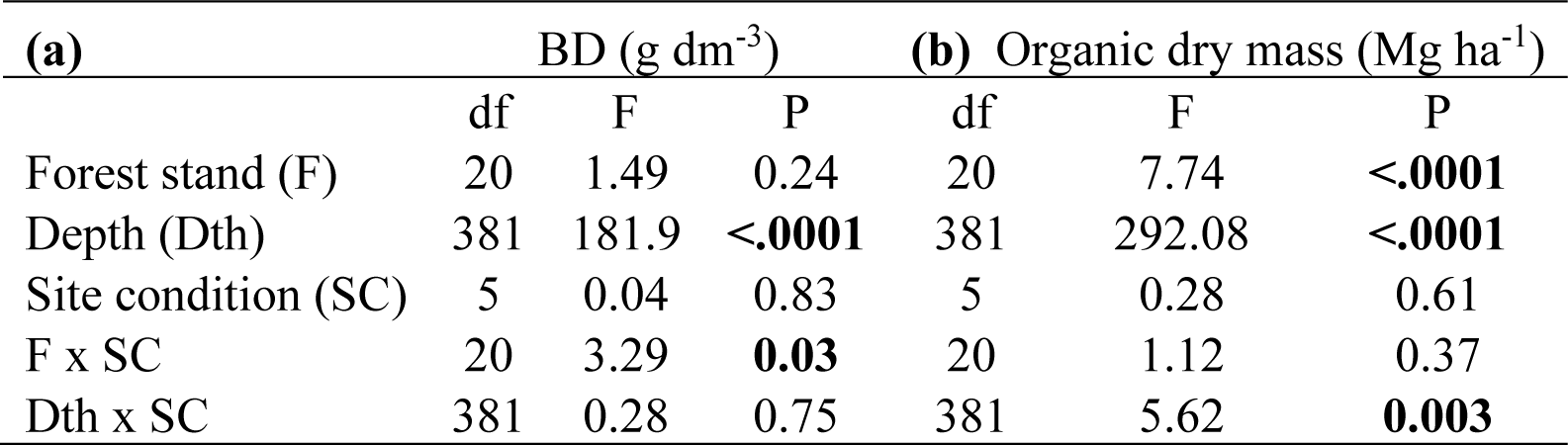
F and P values of the linear mixed effect models on the effect of forest stand type (Be, European beech; D, Douglas fir; S, Norway spruce; SB, Norway spruce/beech; and DB, Douglas fir/beech), site condition (SL, southern sites, loamy soils; and NS, northern sites, sandy soils) and depth [Mineral soil (a) (depth: 0–5, 5–10, and 10–30 cm). Organic layer (b) (layer: Ol, Of and Oh)], on bulk soil density (BD)(g dm^-3^) and organic dry matter (Mg ha^-1^)

The mineral soil bulk density (BD) ranges from 0.8 to 1.4 g cm^-3^. A significant interaction among Forest stand (FS) and Site conditions (SC) was observed (Table 2; p<0.03). At the Southern sites, Douglas fir and Douglas fir/beech showed significant high bulk soil density (p<0.001) compared to beech. Additionally, Douglas fir and its mixture with beech (Douglas fir/beech) showed higher soil densities at loamy soils (southern sites) than at sandy soils (northern sites) (Fig. S3).

### 3.2. C and N concentrations

We used a linear mixed model (LMM) to investigate the relationship between carbon (C) and nitrogen (N) concentrations and the terms Forest stand (F), Depth (Dth), and their interaction across the Site conditions (SC; northern and southern sites). The results showed that Forest stand have a significant effect on C concentration, as well as the Depth and Sites conditions (Table 3a). The highest C concentration was found in the Ol layer and decreased with depth. Furthermore, the effect of Depth varied across Forest stand, with higher carbon concentrations found in the top layer of the organic layer for all forest stand type (Fig S4a).

**Table 3.** F and P values of the linear mixed effect models on the effect of forest stand type (Be, European beech; D, Douglas fir; S, Norway spruce; SB, Norway spruce/beech; and DB, Douglas fir/beech), site condition (SL, southern sites, loamy soils; and NS, northern sites, sandy soils) and depth [(a). Organic layer (layer: Ol, Of and Oh)] and (b) Mineral soil (depth: 0–5, 5–10, and 10–30 cm) on C and N concentrations, and stoichiometric ratios (C:N, C:P and N:P).

| Organic layer |  |  |  |  |  |  |  |  |  |
| --- | --- | --- | --- | --- | --- | --- | --- | --- | --- |
| (a) |  | C (%) |  |  | N (%) |  |  | C:N ratio |  |
|  | df | F | P | df | F | P | DF | F | P |
| Forest stand (F) | 24 | 3.73 | **0.01** | 24 | 1.63 | 0.19 | 24 | 7.69 | **<.0001** |
| Depth (Dth) | 405 | 195.8 | **<.0001** | 405 | 60.16 | **<.0001** | 405 | 297.31 | **<.0001** |
| Site condition (S) | 6 | 11.54 | **0.01** | 6 | 0.1 | 0.75 | 6 | 27.21 | **0.006** |
| F x S | 24 | 2.24 | 0.09 | 24 | 1.14 | 0.35 | 24 | 3.07 | **0.03** |
| Dth x S | 405 | 4.26 | **0.01** | 405 | 0.69 | 0.5 | 405 | 6.34 | **0.001** |
| C:P ratio |  |  |  | N:P ratio |  |  |  |  |  |
|  | df | F | P | df | F | P |  |  |  |
| Forest stand (F) | 24 | 4.15 | **0.01** | 24 | 1.91 | 0.14 |  |  |  |
| Depth (Dth) | 405 | 99.86 | **<.0001** | 405 | 0.23 | 0.79 |  |  |  |
| Site condition (S) | 6 | 17.04 | **0.006** | 6 | 5.18 | 0.06 |  |  |  |
| F x S | 24 | 2.04 | 0.12 | 24 | 1.11 | 0.37 |  |  |  |
| Dth x S | 405 | 37.23 | **<.0001** | 405 | 46.57 | **<.0001** |  |  |  |
| Mineral soil |  |  |  |  |  |  |  |  |  |
| (b) |  | C (%) |  |  | N (%) |  |  | C:N ratio |  |
|  | df | F | P | df | F | P | DF | F | P |
| Forest stand (F) | 23 | 1.46 | 0.24 | 23 | 0,90 | 0.477 | 23 | 2.62 | 0.06 |
| Depth (Dth) | 408 | 351.26 | **<.0001** | 408 | 298.28 | **<.0001** | 408 | 87.54 | **<.0001** |
| Site condition (S) | 6 | 1.73 | 0.23 | 6 | 9.92 | **0.01** | 6 | 37.82 | **0.002** |
| F x S | 23 | 4.18 | **0.01** | 23 | 5.24 | **0.003** | 23 | 0.97 | 0.44 |
| Dth x S | 408 | 2.76 | 0.06 | 408 | 2.39 | 0.09 | 408 | 13.16 | **<.0001** |

In terms of the Forest stand effect, the C concentration was significantly higher under beech and mixed spruce/beech stands compared to pure conifers stands, Douglas fir and spruce. Except at the Oh layer where both conifers and mixed stands had higher concentrations of C than beech stands.

The Depth (Dth) was the only significant term that affected the N concentration (Table 3a), with higher N concentration observed at the Oh layer compared to Of (Fig. S4b).

In terms of C and N concentration on the mineral soil, the interaction term Site conditions (SC) and Forest stand *(SC x F)* was significant (Table 3b). Higher N concentrations were observed at the Southern sites than at the Northern sites, for all forest stand types, expect Douglas fir, where no differences between sites were identified. At the Southern sites, spruce, beech and its mixture (SB) showed higher N concentration than Douglas fir. The same pattern was observed for C concentration (S, SB and B > D). Interestingly, the opposite was observed at the Northern sites, where the highest C concentration was observed under Douglas fir stands.

The stochiometric ratios (C:N, C:P and N:P) were calculated for the organic layer (Table 3a) and the interaction term Forest stand and Site condition shows to be not significant, expect for the C:N ratio, where higher C:N ratio was observed on Douglas fir at the Southern sites (loamy soils) than at the Northern sites (sandy soils) (Fig S6).

### 3.3. C and N stocks

The results of the LMM for C and N stocks revealed significant effects of the terms Forest stand (F) and Depth (Dth) on both variables, with Dth exhibiting a highly significant effect. Meanwhile, the term Site condition (SC) shows no significant impact on C stocks for both, the organic layer and the mineral soil, it shows a notable effect on N stock in the mineral soil (Table 4). Interaction effects, Forest stand and Site condition *(FxSC)*, and Depth and Site condition *(DthxSC)*, were significant for the mineral soil. The results indicated that C stocks were higher in the beech stands at the Southern sites (loamy soils) (23.33 Mg ha^-1^) than in Douglas-fir stands (19.60 Mg ha^-1^). On the contrary, at the Northern sites (sandy soils), Douglas-fir stands had higher C stock (23.46 Mg ha^-1^) than the beech stands (16.29 Mg ha^-1^) (Fig. 1b).

**Figure 1.**
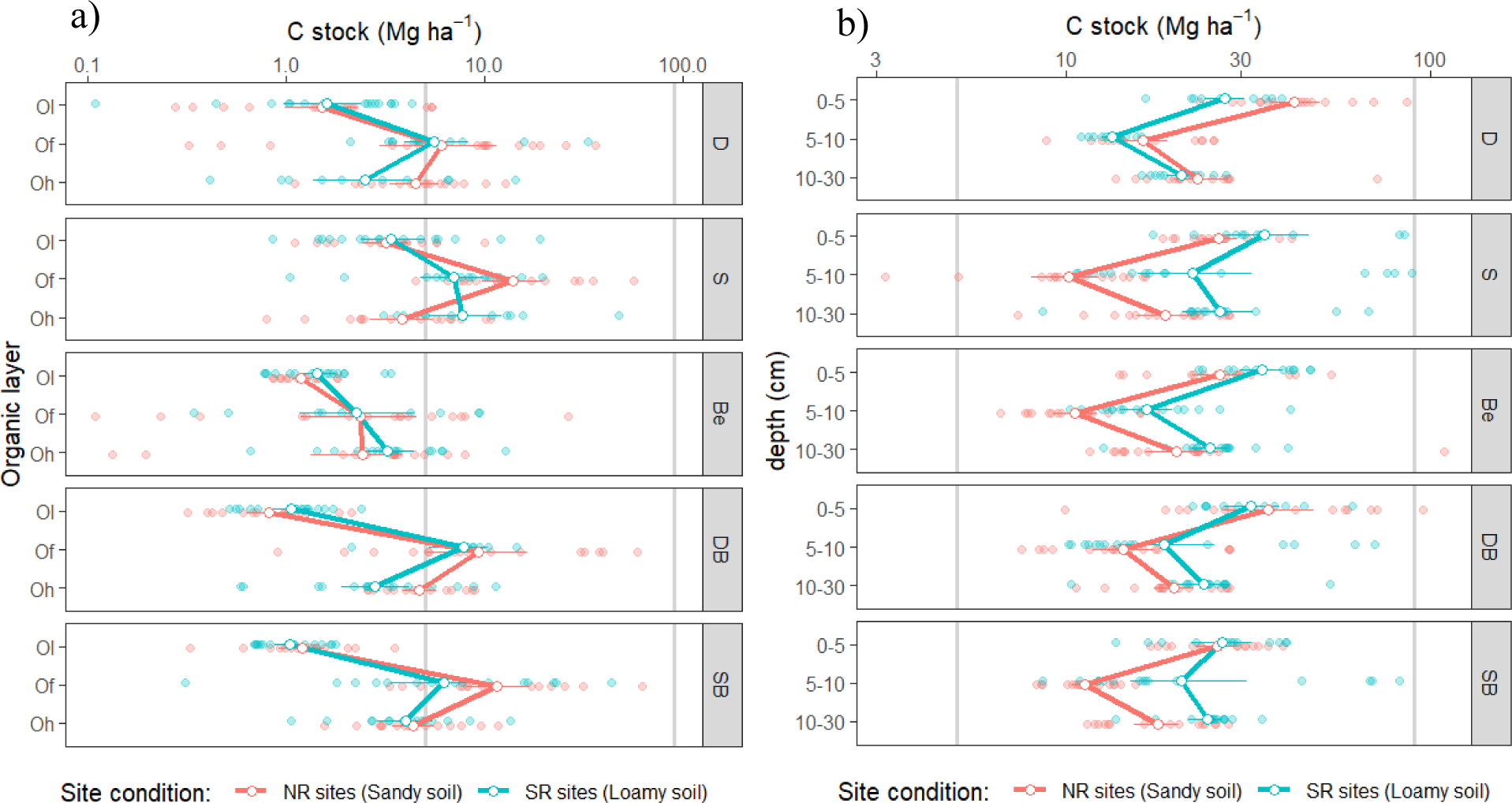
Carbon stocks (Mg ha^-1^) for the organic layer (a) and mineral soil (b). Species (D: Douglas fir, S: spruce; Be: beech; Douglas fir/beech: DB; and, spruce/beech: SB), Site conditions (SR: Southern sites [loamy soils] and NR: Northern sites [sandy soils]) and depth [organic layers: Ol, Of and Oh; mineral soil: 0-5, 5-10 and 10-30 cm]). The points represent means and the horizontal bars the standard errors (n=20).

**Table 4.** F and P values of the linear mixed effect models on the effect of forest stand type (Be, European beech; D, Douglas fir; S, Norway spruce; SB, Norway spruce/beech; and DB, Douglas fir/beech), site condition (SL, southern sites, loamy soils; and NS, northern sites, sandy soils) and depth [(a). Organic layer (layer: Ol, Of and Oh)] and (b) Mineral soil (depth: 0–5, 5–10, and 10–30 cm) on C and N stocks (Mg ha^-1^).

| (a) |  | C stock |  | N stock |  |  |
| --- | --- | --- | --- | --- | --- | --- |
|  | df | F | P | df | F | P |
| Forest stand (F) | 24 | 5.39 | **0.003** | 24 | 5.23 | **0.003** |
| Depth (Dth) | 406 | 126.62 | **<.0001** | 406 | 204.67 | **<.0001** |
| Site condition (S) | 6 | 0.24 | 0.63 | 6 | 0.16 | 0.7 |
| F x S | 24 | 0.49 | 0.73 | 24 | 0.52 | 0.71 |
| Dth x S | 406 | 3.77 | **0.02** | 406 | 2.24 | 0.1 |
| (b) |  | C stock |  | N stock |  |  |
|  | df | F | P | df | F | P |
| Forest stand (F) | 23 | 1.52 | 0.22 | 23 | 0.97 | 0.44 |
| Depth (Dth) | 408 | 232.22 | **<.0001** | 408 | 494.6 | **<.0001** |
| Site condition (S) | 6 | 1.85 | 0.22 | 6 | 9.45 | **0.02** |
| F x S | 23 | 3.8 | **0.01** | 23 | 5.12 | **0.004** |
| Dth x S | 408 | 4.82 | **0.008** | 408 | 7.82 | **<.0001** |

In terms of N stocks, the term Forest stand had a significant effect (p = 0.003), only in the organic layer. Aditionally, the interaction term Forest stand and Site condition *(FxSC)* was significant (p >0.001), only in the mineal soil. The results shows that the effect of forest stand on N stock was attenueted in sandy soils, where Douglas fir has higher N stocks than the spruce stand (Fig. 2b). In contrast, at the loamy soils (southern sites), total N stocks were higher under spruce stands than under Douglas fir stands (Fig. 2b).

**Figure 2.**
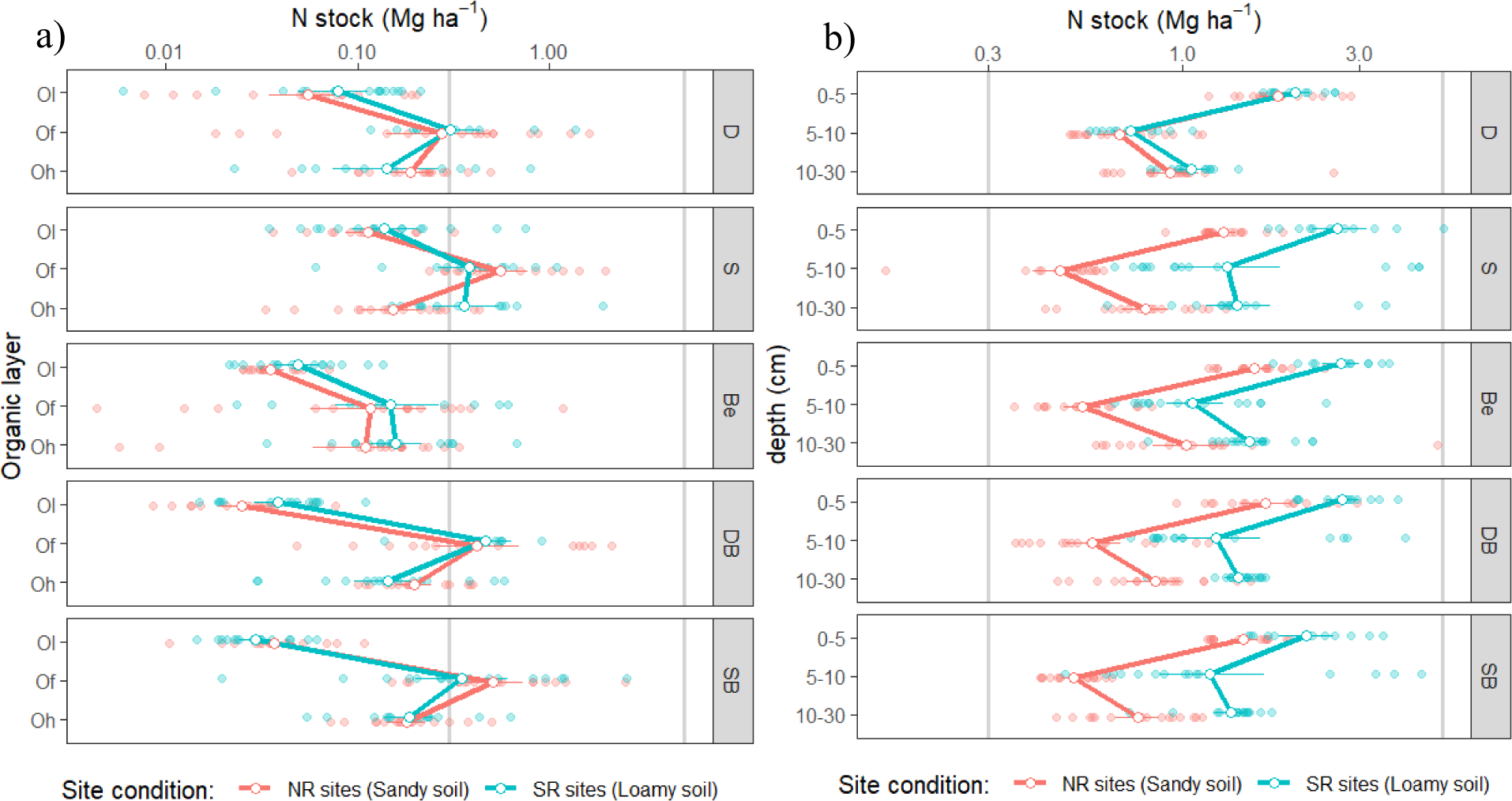
Nitrogen stocks (Mg ha^-1^) for the organic layer (a) and mineral soil (b). Species (D: Douglas fir, S: spruce; Be: beech; Douglas fir/beech: DB; and, spruce/beech: SB), Site conditions (SR: Southern sites [loamy soils] and NR: Northern sites [sandy soils]) and depth [organic layers: Ol, Of and Oh; mineral soil: 0-5, 5-10 and 10-30 cm]). The points represent means and the horizontal bars the standard errors (n=20).

The C stocks under both beech–conifer mixtures were similar to those under the respective pure conifer stands. At the Southern sites, significantly larger C stocks were observed under the mixed spruce/beech stand than Douglas/beech, whereas at the Nouthern sites, the opposite were observed (Douglas/beech > Spruce/beech).

At all sites, N stocks at the organic layer under mixed beech–conifer stands (either Douglas/beech or spruce/beech), significantly exceeded the N stocks in beech stands.

### 3.4. SOC Distribution

The contribution of the organic layer stocks (SOC_Org_) to the total stocks (SOC_t_) was site-dependent. At the Southern sites, the SOC_Org_ contributed between 15% and 22% for Douglas fir and spruce stands, respectively, whereas for beech it was only 8%. The mixture stands showed intermediate results, ranging from 10% (Douglas fir/beech) to 17% (spruce/beech). At the Northern sites, the contribution of SOCOrg to the SOCt stock was strong pronounced.

At the spruce stand, 33% of the SOC stock is allocated to the organic layer, whereas 20% was observed in the Douglas fir stand. The contribution of SOC_org_ to the SOC_t_ for mixed forests ranged between 23% (Douglas/beech) and 30% (spruce/beech) and only 10% for beech stands.

The N allocation corresponded to the SOC vertical distribution. At the Northern sites, the relative contribution ranged from 20–26% for conifers and mixed forest to 11% at the beech stands, whereas at the Southern sites, it ranged from 13–17% for conifers to only 6% under beech forest. At the Northern sites, mixture stands showed a clear contrast: at the spruce/beech stand, the contribution of organic layer N stock to the total N stock was 30%, while at the Douglas/beech stand, it was only 7%.

Overall, under sandy soil conditions, conifers and mixed forests allocated 10% more SOC and N at the organic layer compared to loamy soils, whereas the SOC and N stocks under beech maintained the same proportion (>90%), independent of the site conditions.

### 3.5. Effect of abiotic factors on SOM and N stocks

The effect of abiotic factors like soil pH on soil organic matter (SOM) was investigated using linear mixed models (Fig. 3). The increase of pH caused significant decreased (P<0.001) on SOM at the mineral soil. The increase of 1 unit of pH (3.75 to 4.75) reduced in about 25 % the total SOM. Whereas, the opposite was observed for N stocks, where the increased of pH showed increases in N stocks.

**Figure 3.**
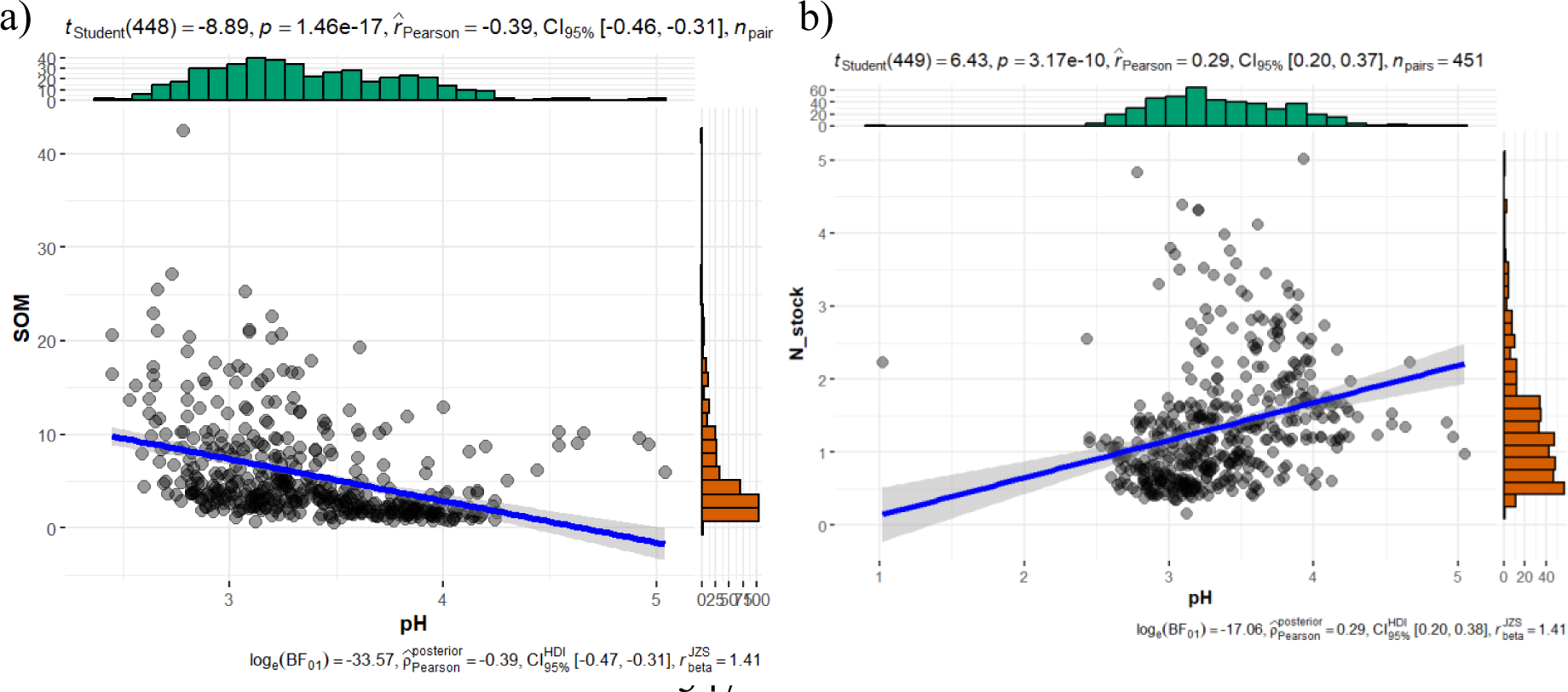
Relationship between the soil organic matter (SOM) (a) and Nitrogen (b) on mineral soil. The grey line denotes a linear mixed-effect model (LMM; P < 0.05) with the predictor variables as fixed effect and species as random intercept terms. The dashed line represents a 95% confidence interval. All response variables (C stock, and N stock) were transformed with natural logarithm, whereas predictor variable (pH) were standardized by subtracting the mean and dividing by the standard deviation.

## 4. Discussion

### 4.1. Tree species effects

We observed significant effects of Forest stand types on the organic C and N stocks as well as their vertical distribution across all investigated sites. Overall, the C and N stocks of the organic layer were significantly higher under conifer forests (Douglas fir and spruce) than under beech (Fig. 1). At the mineral soil, the effects of forest stand type were site-dependent. At the Southern sites (loamy soils), beech and spruce stands accumulated higher C and N stocks than Douglas fir forests, confirming earlier results of Antisari et al. (2015), where smallest SOC stocks of the organic layer under beech stand are accompanied by enhanced SOC stock in the mineral topsoil. In contrast, at the Northern sites (sandy soil), the beech stands showed the smallest C stocks at both layers, organic and mineral soil.

According to Neumann et al. (2018), litter of broadleaves tends to decompose faster compared to conifer litter. This can be attributed to lower lignin and phenol concentrations, which promote rapid decomposition rates and efficient accumulation of mineral-associated organic matter, as reported by Rasse et al. (2005). In our study, we observed a carbon stock shift from the organic layer to the mineral soil in the beech forest under loamy soil conditions, supporting the findings of previous research (Achilles et al., 2021). The higher allocation of C in the mineral soil than in the organic layer under beech forests has also been reported in common garden studies (Vesterdal et al., 2013), when compared to conifers. These differences in the vertical distribution have been attributed to the associated community of macrofauna species in forests dominated by beech (Achilles et al., 2021). Endogenic earthworm had the potential for slowly altering SOC pools in the upper mineral soil of beech forests (Heinze et al., 2021), feeding and translocating litter of the forest floor and incorporating organic matter into the topsoil (Scheu et al., 2002). These earthworms ingested large amounts of topsoil organic matter and translocated SOC into deeper horizons by bioturbation (Frouz et al., 2009). The depth distribution of soil organic carbon, with significant lower C in forest floor layers (Ol-layer) and a noticeably higher concentration in the topsoil (Of - Oh layers and 0-5 cm) (Fig. 1), point to carbon translocation processes in beech forests. This translocation from the forest floor into the topsoil may build up stable carbon pools and contributes to carbon sequestration (Vesterdal et al., 2013).

Therefore, the quality of the soil organic matter (SOM) might be significantly different under coniferous and deciduous trees (Jaffrain et al., 2007).The O_H_ layer under conifers results in a more recalcitrant and hydrophobic composition of the humus than that of O_H_ layer at beech stand (Thomas et al., 2014), therefore, slowing down the C turnover of the SOM, which leads to a greater accumulation of carbon in the transition layers, such as the Of and Oh layers, and further to the top mineral soil, as observed in our study under both conifers stands.

Besides the litter quality, rooting patterns and microbial activity among beech, Douglas fir and Norway spruce might change the C inputs between tree species. Conifer roots contain lower lignin concentrations than roots of beech (Eckhart et al., 2019; Thomas et al., 2014), which might lead to a lower longevity and a faster decomposition. Additionally, fine roots necromass deliver a considerable amount of organic material (Cremer et al., 2016; Dawud et al., 2017) enhancing the root-derived SOM. Simultaneous studies to the one presented here on the turnover of fine root biomasses carried out by Lwila et al. (2021) have shown that beech forests showed higher fine roots biomass than Douglas fir at the sandy soils, whereas the fine root necromass was higher for Douglas fir compared to beech. Additionally, high microbial activity at the upper mineral soil is reported in the same study area (Lu and Scheu, 2021), suggested higher rhizosphere carbon inputs at the beech forest stand compared to Douglas fir, but also high root-derived SOM at the Douglas fir stand.

### 4.2. Mixed forest effect

Independent of the site conditions, the pure stands of beech showed relatively small total SOC stocks, whereas the pure conifer sites enhanced total SOC and N stocks, accumulated largely in the organic layer. Furthermore, the admixture of conifers to beech enlarged the total SOC and N storage only at the Northern sites, whereas at the Southern sites, both applied mixtures of conifers to beech stands presented similar storages compared to pure beech. Overall, the carbon (C) stocks in the Oh layer at the Southern sites, as well as SOC and N stocks, were generally similar between mixed beech-conifer stands and their respective pure stands. The SOC stocks at both mixed stands at the Southern sites resembled those of the respective beech stands and approached the levels observed in conifer stands at the Northern sites.

The admixture of conifers into beech stands in nutrient-poor soil conditions, particularly observed at the Northern sites, had significant effects on SOC and N stocks, which aligns with the results reported by Dawud et al. (2017). Under these soil conditions (nutrient-poor soils), both stands, the mixture of Douglas fir and beech and pure Douglas fir stands exhibited higher SOC stocks in the humus layer (Oh) and the 5-30 cm soil depth. This can be attributed to the competitive advantage of beech fine roots, which can access deeper soil layers and exploit areas less occupied by competing species. In contrast, Douglas fir fine roots tend to be restricted to the topsoil (15-30 cm), resulting in a shift in the distribution of beech fine roots towards the subsoil. This phenomenon has been reported in previous studies (Hendriks and Bianchi, 1995; Schmid and Kazda, 2002). These observed shifts of root vertical distribution are known as below-ground complementarity effects, e.g., through vertical segregation of roots of different species, which exploit soil at different depths allowing for reduced competition (Loreau M, 2001). It has been suggested as a mechanism by which species mixtures stands store more C in deeper soil layers than monocultures (Bauhus et al., 2009; Forrester et al., 2010). Additionally, the admixture of beech leaf litter to the more recalcitrant needle litter induced a faster litter mass loss and consequently shift of SOC stock from Ol to Oh layer. We observed similar litter decomposition rate for mixed stands and pure beech, and higher than pure conifers (Douglas fir and spruce), suggesting similar decomposability of litter in the mixed forest and in the beech forest (not published).

However, sequestration of C in soils is an important ecosystem service provided by forests (Pretzsch et al., 2017). Due to the potential of elevated SOC storages in mixed-species forests compared to pure stands and to shift the C into stable pools, translocating carbon from the forest floor to more stable carbon pools such as the humus layer and upper mineral soil, enhance the capacity of mixed forests to provide ecosystem services, such as improving soil biodiversity and nutrient availability. Furthermore, it may contribute to the resilience of the forest ecosystem in adapting to future climate change scenarios.

### 4.3. Abiotic effects

The persistence of soil organic Carbon (SOC) is largely due to complex interactions between SOC and its environment, such as reactive mineral surfaces, climate, water availability and soil acidity (Leifeld et al., 2013; Mikutta et al., 2009; Zhou et al., 2019). The most important factor in SOC stabilization at sites with high clay contents, as partly observed in our study, is probably the association with soil minerals, irrespective of forest type (Fierer and Jackson, 2006; Meier et al., 2020). Indeed, enhanced clay and silt content increased the SOC in the mineral soil at our investigated sites. Loamy soils are known to buffer influences by tree species more strongly than sandy soils (Meier and Leuschner, 2010), and this would explain why we see clear effects of forest stand type at the Northern sites dominated by sandy soils than at the Southern sites, dominated by loamy soils.

Therefore, differences in soil pH often go along with differences in soil mineralogy as well and the latter exerts control on the stabilization of mineral associated organic matter (Meier et al., 2020) and microbiota community. In our study, we found that the increase of pH caused significant decreased (P<0.001) on soil organic matter (SOM) at mineral soil (Fig. 3). Thus, the accumulation of organic matter in soils is influenced by relatively low pH levels, which can reduce the decomposition of SOM, as highlighted by Meier et al. (2020). This, in turns, leads to noticeable effects of forest stand type on organic matter accumulation at the Northern sites, where lower pH is observed, as observed in our study. Interestingly, Meier et al. (2020) also suggested that soil acidity is associated with increased rates of root exudation in beech forests, particularly in the topsoil of glacial sandy soils found at the Northern sites. The increased root exudation can be seen as an adaptation of the trees to low nutrient availability, a finding consistent with our study (Foltran et al., 2023). In these conditions, the majority of nutrient uptake occurs in the topsoil, which is enriched with organic material, further explaining the adaptability and plasticity of beech forests in response to varying nutrient availability.

## 5. Conclusion

Site dependent effects of tree species (European beech, Douglas fir and Norway spruce) on SOC and N stocks as well as on OC and N concentration were observed. The SOC and N stocks generally were smallest in pure beech stands compared with Douglas fir and Norway spruce. The SOC and N stocks in mixed stands of beech with Douglas fir or Norway spruce are generally between those of the respective pure stands. However, the results indicate that the effects of admixture of conifers into beech stands are site-dependent. At the Northern sites, the adaptability of Douglas fir under dry and nutrient-poor site conditions, and combined with high plasticity of beech, seems to promote a favorable effect on total SOC and N, enhancing its storages as observed at pure Douglas fir but also in the mixture Douglas fir/beech stands. In contrast, at the Southern sites, spruce/beech stands showed higher SOC and N stocks than mixed Douglas/beech stands. Additionally, the potential shift of carbon into more stable pools (shifts from the Ol layer to the Oh layer), emphasizes the capacity of mixed forest to provide valuable ecosystem services, enhancing C sequestration, meanwhile muting the risk of unintended losses.

## Author contributions

All authors contributed to the study conception and design. Material preparation, data collection and analysis were performed by EF. The first draft of the manuscript was written by EF and NL commented on previous versions of the manuscript. All authors read and approved the final manuscript.

## Acknowledgements

The study was conducted as part of the Research Training Group 2300 funded by the German research funding organization (Deutsche Forschungsgemeinschaft – DFG). We gratefully acknowledge the administrative support by Serena Müller and the indispensable help of Julian Meyer and Dirk Böttger during soil sampling. Furthermore, we thank Sylvia Bondzio, Karin Schmidt for their valuable advice during laboratory work.

## Conflicts of Interest

The authors declare no conflict of interest.

Declaration of generative AI and AI-assisted technologies in the writing process

During the preparation of this work the authors used the tool Grammarly in order to correct the english grammar. After using this tool, the authors reviewed and edited the content as needed and takes full responsibility for the content of the publication.

## References

Achilles, F., Tischer, A., Bernhardt-r, M., Heinze, M., Reinhardt, F., Makeschin, F., Michalzik, B., 2021. European beech leads to more bioactive humus forms but stronger mineral soil acidification as Norway spruce and Scots pine – Results of a repeated site assessment after 63 and 82 years of forest conversion in Central Germany. Forest Ecology and Management 483. 10.1016/j.foreco.2020.118769

Adams, W.A., 1973. THE EFFECT OF ORGANIC MATTER ON THE BULK AND TRUE DENSITIES OF SOME UNCULTIVATED PODZOLIC SOILS. Journal of Soil Science 24, 10–17. 10.1111/j.1365-2389.1973.tb00737.x

Ammer, C., 2019. Diversity and forest productivity in a changing climate. New Phytologist 221, 50–66. 10.1111/nph.15263

Angst, G., Messinger, J., Greiner, M., Häusler, W., Hertel, D., Kirfel, K., Kögel-Knabner, I., Leuschner, C., Rethemeyer, J., Mueller, C.W., 2018. Soil organic carbon stocks in topsoil and subsoil controlled by parent material, carbon input in the rhizosphere, and microbial-derived compounds. Soil Biology and Biochemistry 122, 19–30. 10.1016/J.SOILBIO.2018.03.026

Antisari, L.V., Falsone, G., Carbone, S., Marinari, S., Vianello, G., 2015. Douglas-fir reforestation in North Apennine (Italy): Performance on soil carbon sequestration, nutrients stock and microbial activity. Applied Soil Ecology 86, 82–90. 10.1016/j.apsoil.2014.09.009

Bauhus, J., Puettmann, K., Messier, C., 2009. Silviculture for old-growth attributes. Forest Ecology and Management 258, 525–537. 10.1016/j.foreco.2009.01.053

Bolte, A., Villanueva, I., 2006. Interspecific competition impacts on the morphology and distribution of fine roots in European beech (Fagus sylvatica L.) and Norway spruce (Picea abies (L.) karst.). European Journal of Forest Research 125, 15–26. 10.1007/s10342-005-0075-5

Cepáková, Š., Tošner, Z., Frouz, J., 2016. The effect of tree species on seasonal fluctuations in water-soluble and hot water-extractable organic matter at post-mining sites. Geoderma 275, 19–27. 10.1016/j.geoderma.2016.04.006

Chen, Y., Huang, Y., Sun, W., 2017. Using Organic Matter and pH to Estimate the Bulk Density of Afforested/Reforested Soils in Northwest and Northeast China. Pedosphere 27, 890–900. 10.1016/S1002-0160(17)60372-2

Cools, N., Vesterdal, L., De Vos, B., Vanguelova, E., Hansen, K., 2014. Tree species is the major factor explaining C: N ratios in European forest soils. Forest Ecology and Management 311, 3–16. 10.1016/j.foreco.2013.06.047

Cremer, M., Kern, N.V., Prietzel, J., 2016. Soil organic carbon and nitrogen stocks under pure and mixed stands of European beech, Douglas fir and Norway spruce. Forest Ecology and Management 367, 30–40. 10.1016/j.foreco.2016.02.020

Dawud, S.M., Vesterdal, L., Raulund-Rasmussen, K., 2017. Mixed-species effects on soil C and N stocks, C/N ratio and pH using a transboundary approach in adjacent common garden douglas-fir and beech stands. Forests 8. 10.3390/f8040095

Dobor, L., Hlásny, T., Rammer, W., Zimová, S., Barka, I., Seidl, R., 2020. Spatial configuration matters when removing windfelled trees to manage bark beetle disturbances in Central European forest landscapes. Journal of Environmental Management 254, 109792. 10.1016/J.JENVMAN.2019.109792

Eckhart, T., Pötzelsberger, E., Koeck, R., Thom, D., Lair, G.J., van Loo, M., Hasenauer, H., 2019. Forest stand productivity derived from site conditions: an assessment of old Douglas-fir stands (Pseudotsuga menziesii (Mirb.) Franco var. menziesii) in Central Europe. Annals of Forest Science 76. 10.1007/s13595-019-0805-3

FAO, 2014. World reference base for soil resources 2014. International soil classification system for naming soils and creating legends for soil maps, World Soil Resources Reports No. 106.

Fierer, N., Jackson, R.B., 2006. The diversity and biogeography of soil bacterial communities. Proceedings of the National Academy of Sciences of the United States of America 103, 626–631. 10.1073/pnas.0507535103

Foltran, E.C., Ammer, C., Lamersdorf, N., 2023. Do admixed conifers change soil nutrient conditions of European beech stands? Soil Research 61, 647–662.

Forrester, D.I., Medhurst, J.L., Wood, M., Beadle, C.L., Valencia, J.C., 2010. Growth and physiological responses to silviculture for producing solid-wood products from Eucalyptus plantations: An Australian perspective. Forest Ecology and Management 259, 1819–1835. 10.1016/j.foreco.2009.08.029

Frouz, J., Pižl, V., Cienciala, E., Kalčík, J., 2009. Carbon Storage in Post-Mining Forest Soil, the Role of Tree Biomass and Soil Bioturbation. Biogeochemistry 94, 111–121.

Glatthorn, J., 2021. A spatially explicit index for tree species or trait diversity at neighborhood and stand level. Ecological Indicators 130, 108073. 10.1016/j.ecolind.2021.108073

Heinze, M., Achilles, F., Tischer, A., Bernhardt-r, M., Reinhardt, F., Makeschin, F., Michalzik, B., 2021. European beech leads to more bioactive humus forms but stronger mineral soil acidification as Norway spruce and Scots pine – Results of a repeated site assessment after 63 and 82 years of forest conversion in Central Germany. Forest Ecology and Management 483. 10.1016/j.foreco.2020.118769

Hendriks, C.M.A., Bianchi, F.J.J.A., 1995. Root density and root biomass in pure and mixed forest stands of Douglas-fir and Beech. Netherlands Journal of Agricultural Science 43, 321–331. 10.18174/njas.v43i3.570

Hlásny, T., Turčáni, M., 2013. Persisting bark beetle outbreak indicates the unsustainability of secondary Norway spruce forests: case study from Central Europe. Annals of Forest Science 70, 481–491. 10.1007/s13595-013-0279-7

Höhle, J., Bielefeldt, J., Dühnelt, P., König, N., Ziche, D., 2018. Bodenzustandserhebung im Wald – Dokumentation und Harmonisierung der Methoden. Thünen Working Paper 97.

Jaffrain, J., Gérard, F., Meyer, M., Ranger, J., 2007. Assessing the Quality of Dissolved Organic Matter in Forest Soils Using Ultraviolet Absorption Spectrophotometry. Soil Science Society of America Journal 71, 1851–1858. 10.2136/sssaj2006.0202

Jandl, R., Lindner, M., Vesterdal, L., Bauwens, B., Baritz, R., Hagedorn, F., Johnson, D.W., Minkkinen, K., Byrne, K.A., 2007. How strongly can forest management influence soil carbon sequestration? Geoderma 137, 253–268. 10.1016/J.GEODERMA.2006.09.003

Kölling, C., Knoke, T., Schall, P., Ammer, C., 2009. Cultivation of Norway spruce (Picea abies (L.) Karst.) in Germany: considerations on risk against the background of climate change (original title in German: Überlegungen zum Risiko des Fichtenanbaus in Deutschland vor dem Hintergrund des Klimawandels). Forstarchiv 80, 42–54.

Kölling, C., Zimmermann, L., 2007. Die Anfälligkeit der Wälder Deutschlands gegenüber dem Klimawandel 67, 259–268.

König, Nils., Blum, U., Symossek, F., Bussian, B., Furtmann, K., Gartner, A., Groeticke, K., Gutwasser, F., Hoehle, J., Hauenstein, M., Kiesling, G., Klingernberg, U., Klinger, T., Nack, T., Stahn, M., Trefz-Malcher, G., Wies, K., 2014. Handbuch Forstliche Analytik: Eine Loseblatt-Sammlung der Analysemethoden im Forstbereich. 568.

Krishna, M.P., Mohan, M., 2017. Litter decomposition in forest ecosystems: a review. Energy, Ecology and Environment 2, 236–249. 10.1007/s40974-017-0064-9

Leifeld, J., Bassin, S., Conen, F., Hajdas, I., Egli, M., Fuhrer, J., 2013. Control of soil pH on turnover of belowground organic matter in subalpine grassland. Biogeochemistry 112, 59–69. 10.1007/s10533-011-9689-5

Loreau M, H.A., 2001. Partitioning selection and complementarity in biodiversity experiments. Nature 412:72–76. Nature 412, 72–76.

Lu, J.-Z., Scheu, S., 2021. Response of soil microbial communities to mixed beech-conifer forests varies with site conditions. Soil Biology and Biochemistry 155, 1–29. 10.1101/2020.07.21.213900

Lwila, A.S., Mund, M., Ammer, C., Glatthorn, J., 2021. Site conditions more than species identity drive fine root biomass, morphology and spatial distribution in temperate pure and mixed forests. Forest Ecology and Management 499. 10.1016/j.foreco.2021.119581

Meier, I., Leuschner, C., 2010. Variation of soil and biomass carbon pools in beech forests across a precipitation gradient. Global Change Biology 16, 1035–1045. 10.1111/j.1365-2486.2009.02074.x

Meier, I.C., Tückmantel, T., Heitkötter, J., Müller, K., Preusser, S., Wrobel, T.J., Kandeler, E., Marschner, B., Leuschner, C., 2020. Root exudation of mature beech forests across a nutrient availability gradient: the role of root morphology and fungal activity. New Phytologist 226, 583–594. 10.1111/nph.16389

Mikutta, R., Schaumann, G.E., Gildemeister, D., Bonneville, S., Kramer, M.G., Chorover, J., Chadwick, O.A., Guggenberger, G., 2009. Biogeochemistry of mineral-organic associations across a long-term mineralogical soil gradient (0.3-4100 kyr), Hawaiian Islands. Geochimica et Cosmochimica Acta 73, 2034–2060. 10.1016/j.gca.2008.12.028

Mueller, K.E., Eissenstat, D.M., Hobbie, S.E., Oleksyn, J., Jagodzinski, A.M., Reich, P.B., Chadwick, O.A., Chorover, J., 2012. Tree species effects on coupled cycles of carbon, nitrogen, and acidity in mineral soils at a common garden experiment. Biogeochemistry 111, 601–614. 10.1007/s10533-011-9695-7

Neumann, G., Martinoia, E., 2002. Cluster roots – an underground adaptation for survival in extreme environments 7, 162–167.

Neumann, M., Ukonmaanaho, L., Johnson, J., Benham, S., Vesterdal, L., Novotný, R., Verstraeten, A., Lundin, L., Thimonier, A., Michopoulos, P., Hasenauer, H., 2018. Quantifying Carbon and Nutrient Input From Litterfall in European Forests Using Field Observations and Modeling. Global Biogeochemical Cycles 32, 784–798. 10.1029/2017GB005825

Oulehle, F., Hofmeister, J., Hruška, J., 2007. Modeling of the long-term effect of tree species (Norway spruce and European beech) on soil acidification in the Ore Mountains. Ecological Modelling 204, 359–371. 10.1016/j.ecolmodel.2007.01.012

Patil, I., 2021. Visualizations with statistical details: The ‘ggstatsplot’ approach. Journal of Open Source Software 6, 3167.

Piovia-Scott, J., Yang, L.H., Wright, A.N., Spiller, D.A., Schoener, T.W., 2019. Pulsed seaweed subsidies drive sequential shifts in the effects of lizard predators on island food webs. Ecology Letters 22, 1850–1859. 10.1111/ele.13377

Pretzsch, H., Forrester, D.I., Bauhus, J., 2017. Mixed-Species Forests, 1st ed. Springer Berlin Heidelberg, Berlin, Germany. 10.1007/978-3-662-54553-9

Rasmussen, C., Heckman, K., Wieder, W.R., Keiluweit, M., Lawrence, C.R., Berhe, A.A., Blankinship, J.C., Crow, S.E., Druhan, J.L., Hicks Pries, C.E., Marin-Spiotta, E., Plante, A.F., Schädel, C., Schimel, J.P., Sierra, C.A., Thompson, A., Wagai, R., 2018. Beyond clay: towards an improved set of variables for predicting soil organic matter content. Biogeochemistry 137, 297–306. 10.1007/s10533-018-0424-3

Rasse, D.P., Rumpel, C., Dignac, M.F., 2005. Is soil carbon mostly root carbon? Mechanisms for a specific stabilisation. Plant and Soil 269, 341–356. 10.1007/s11104-004-0907-y

Scheu, S., Schlitt, N., Tiunov, A.V., Newington, J.E., Jones, H.T., 2002. Effects of the presence and community composition of earthworms on microbial community functioning. Oecologia 133, 254–260. 10.1007/s00442-002-1023-4

Schmid, I., Kazda, M., 2002. Root distribution of Norway spruce in monospecific and mixed stands on different soils. Forest Ecology and Management 159, 37–47. 10.1016/S0378-1127(01)00708-3

Thomas, F.M., Molitor, F., Werner, W., 2014. Lignin and cellulose concentrations in roots of Douglas fir and European beech of different diameter classes and soil depths. Trees - Structure and Function 28, 309–315. 10.1007/s00468-013-0937-2

Vesterdal, L., Clarke, N., Sigurdsson, B.D., Gundersen, P., 2013. Do tree species influence soil carbon stocks in temperate and boreal forests? Forest Ecology and Management 309, 4–18. 10.1016/j.foreco.2013.01.017

Vesterdal, L., Raulund-Rasmussen, K., 1998. Forest floor chemistry under seven tree species along a soil fertility gradient. Canadian Journal of Forest Research 28, 1636–1647. 10.1139/cjfr-28-11-1636

Vesterdal, L., Ritter, E., Gundersen, P., 2002. Change in soil organic carbon following afforestation of former arable land. Forest Ecology and Management 169, 137–147. 10.1016/S0378-1127(02)00304-3

Vesterdal, L., Schmidt, I.K., Callesen, I., Nilsson, L.O., Gundersen, P., 2008. Carbon and nitrogen in forest floor and mineral soil under six common European tree species. Forest Ecology and Management 255, 35–48. 10.1016/j.foreco.2007.08.015

Zhou, W., Han, G., Liu, M., Li, X., 2019. Effects of soil pH and texture on soil carbon and nitrogen in soil profiles under different land uses in Mun River Basin, Northeast Thailand. PeerJ 2019. 10.7717/peerj.7880

