## Supplementary material for "Tree species identity drives soil Carbon and Nitrogen stocks in nutrient-poor sites": Suplemental figures

**Supporting information**

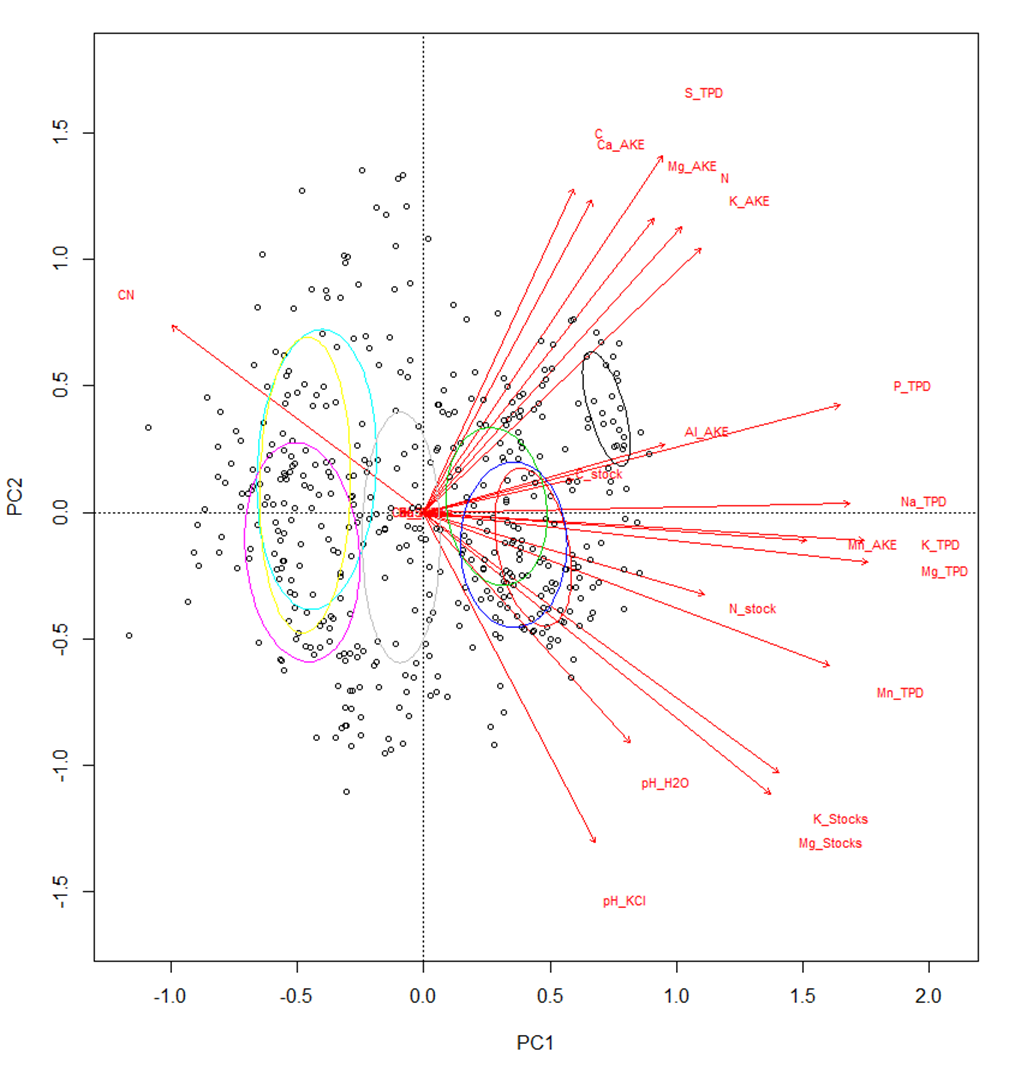

Fig S1. PCA analysis considers all soil variables from mineral soil datasets (Foltran et al. 2023). The cluster shows that all plots (1-8), therefore, clustering into 2 distinctive groups, nutrient-rich sites (loamy soils) and nutrient-poor sites (sandy soils).

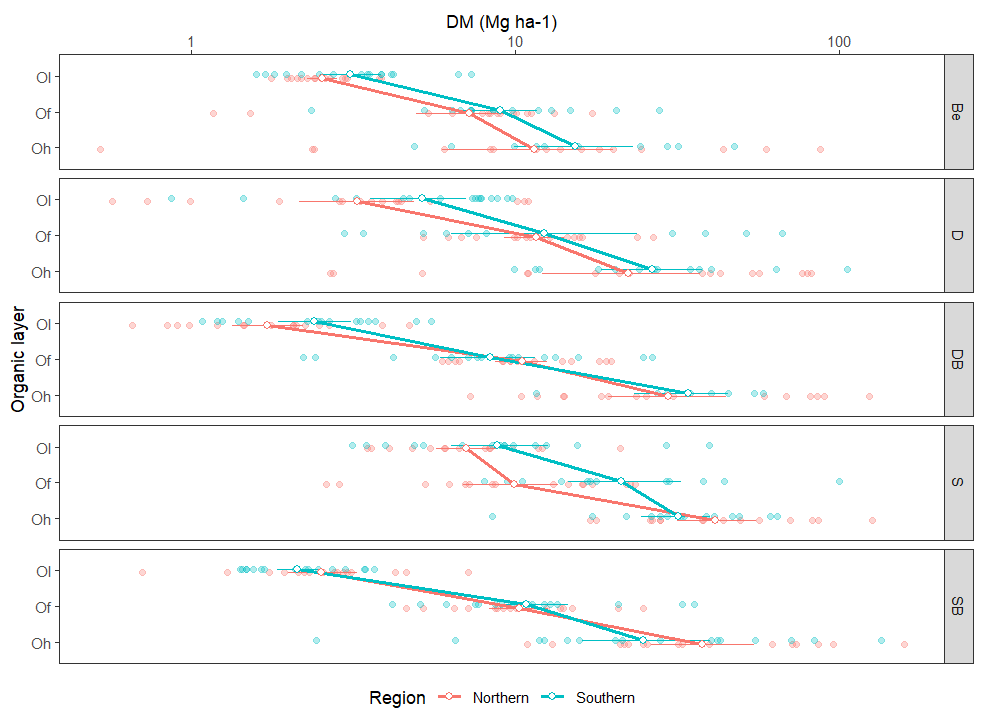

Fig.S2. Organic dry mass (Mg ha^-1^) for the organic layer. Species (D: Douglas fir, S: spruce; Be: beech; Douglas fir/beech: DB; and, spruce/beech: SB), Site conditions (SR: Southern sites [loamy soils] and NR: Northern sites [sandy soils]) and depth (organic layers: Ol, Of and Oh). The points represent means and the horizontal bars the standard errors (n=20).

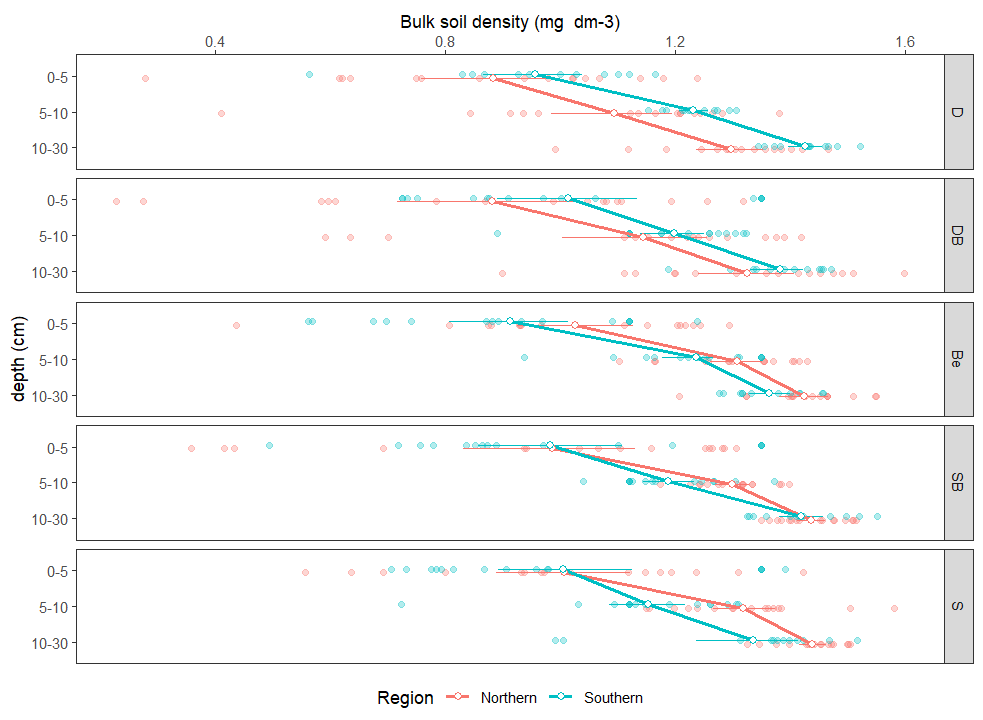

Fig S3. Soil bulk density (mg dm-³) for the mineral soil. Species (D: Douglas fir, S: spruce; Be: beech; Douglas fir/beech: DB; and, spruce/beech: SB), Site conditions (SR: Southern sites [loamy soils] and NR: Northern sites [sandy soils]) and depth (mineral soil: 0-5, 5-10 and 10-30 cm). The points represent means and the horizontal bars the standard errors (n=20).

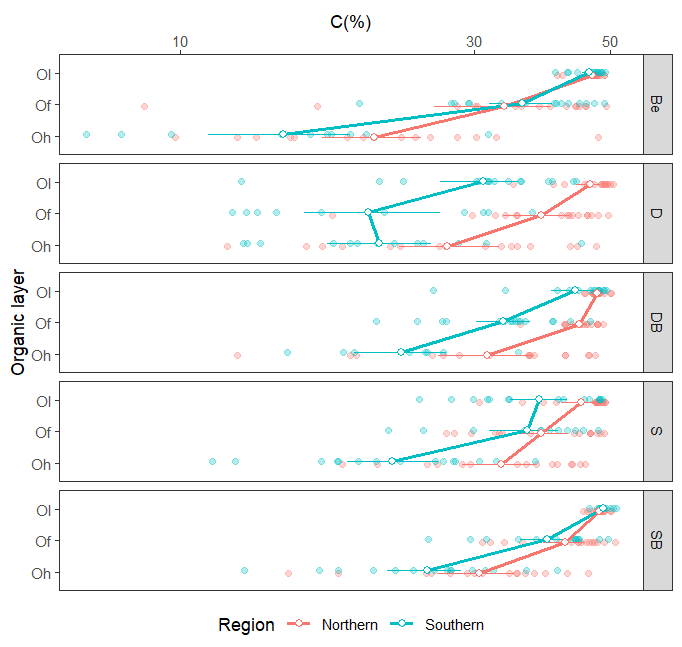

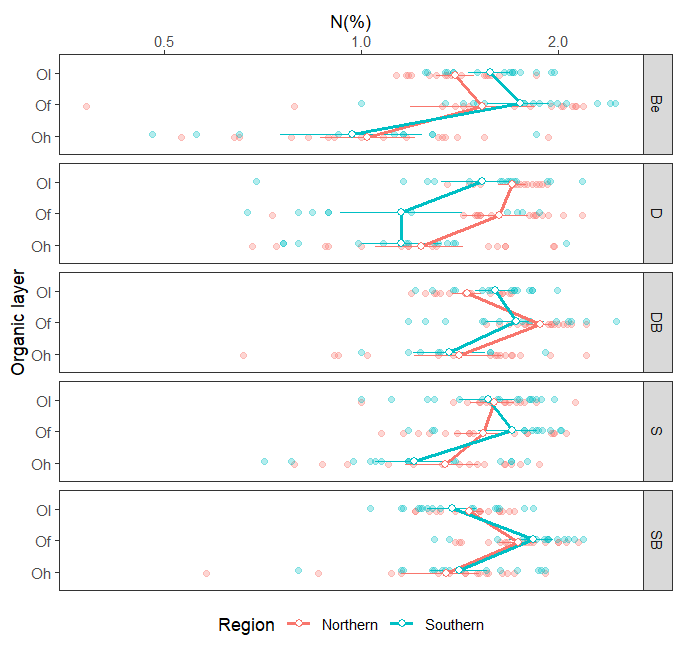

Fig S4. Carbon (%) (a) and Nitrogen (%) (b). Species (D: Douglas fir, S: spruce; Be: beech; Douglas fir/beech: DB; and, spruce/beech: SB), Site conditions (SR: Southern sites [loamy soils] and NR: Northern sites [sandy soils]) and depth (organic layers: Ol, Of and Oh). The points represent means and the horizontal bars the standard errors (n=20).

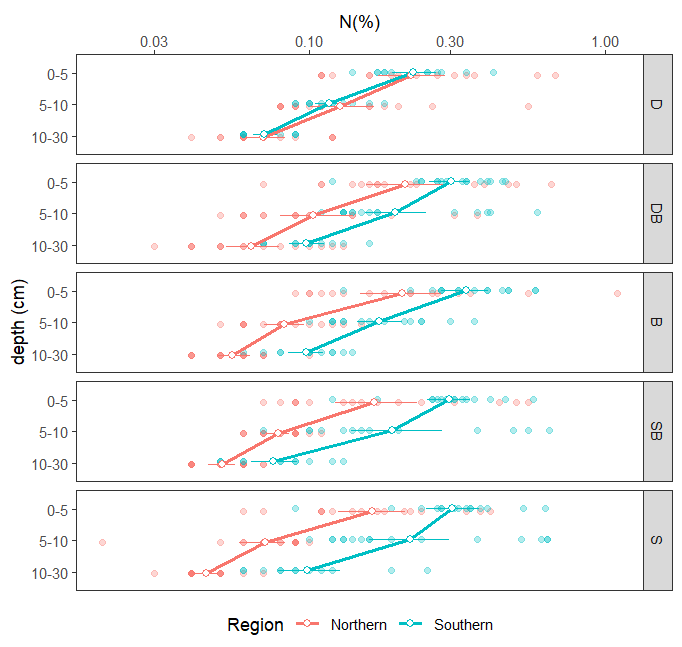

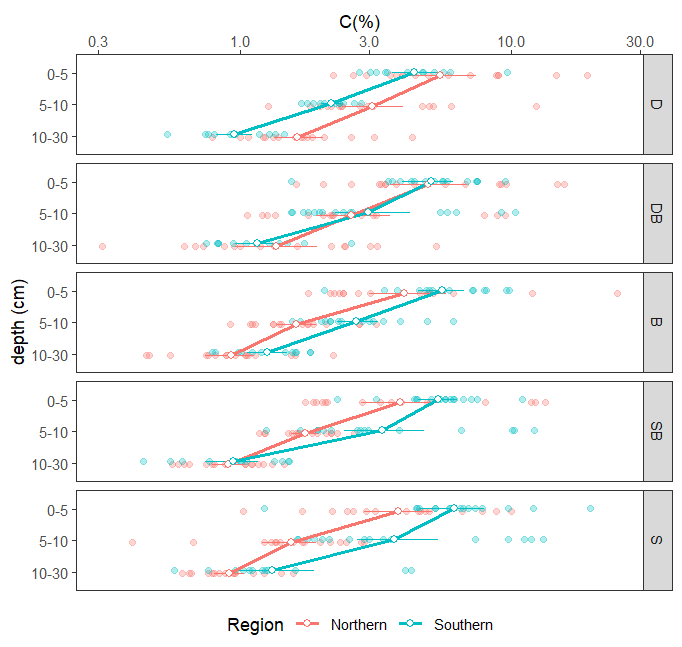

Fig S5. Carbon (%) (a) and Nitrogen (%) (b). Species (D: Douglas fir, S: spruce; Be: beech; Douglas fir/beech: DB; and, spruce/beech: SB), Site conditions (SR: Southern sites [loamy soils] and NR: Northern sites [sandy soils]) and depth (mineral soil: 0-5, 5-10 and 10-30 cm). The points represent means and the horizontal bars the standard errors (n=20).

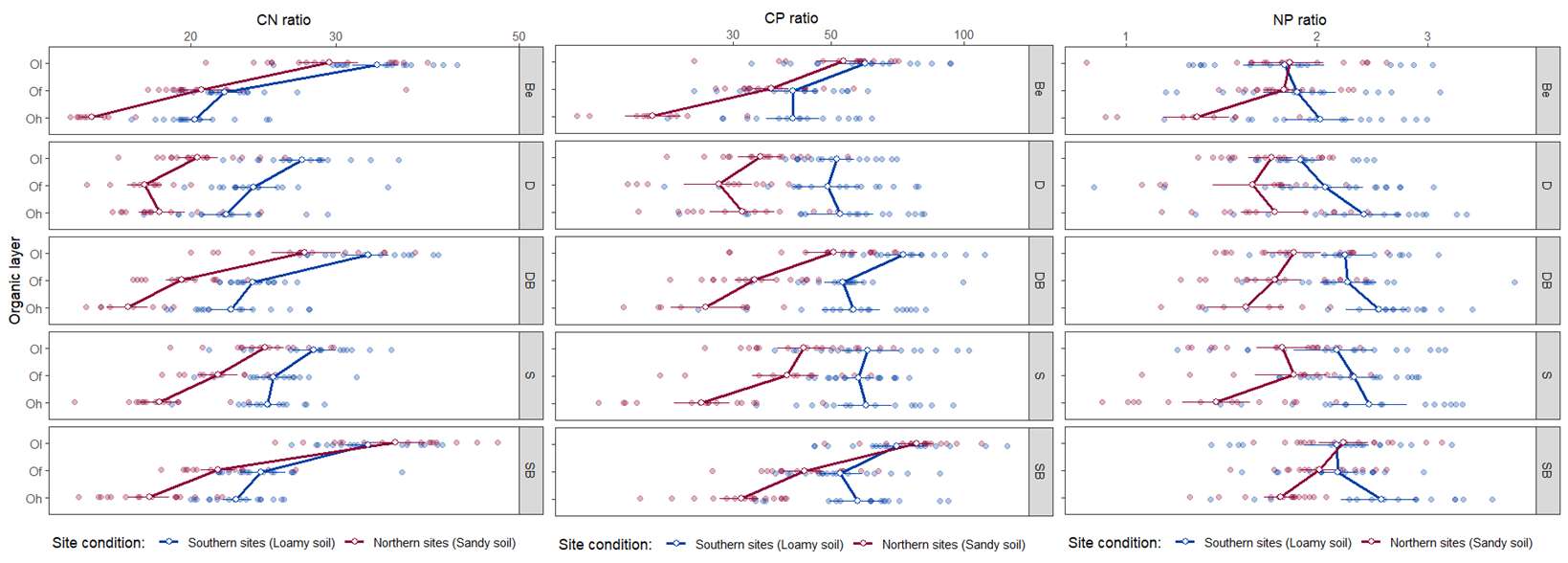

Fig S6. Stoichiometric ratios (CN, CP and NP) for the organic layers. Species (D: Douglas fir, S: spruce; Be: beech; Douglas fir/beech: DB; and, spruce/beech: SB), Site conditions (SR: Southern sites [loamy soils] and NR: Northern sites [sandy soils]) and depth (organic layers: Ol, Of and Oh). The points represent means and the horizontal bars the standard errors (n=20).

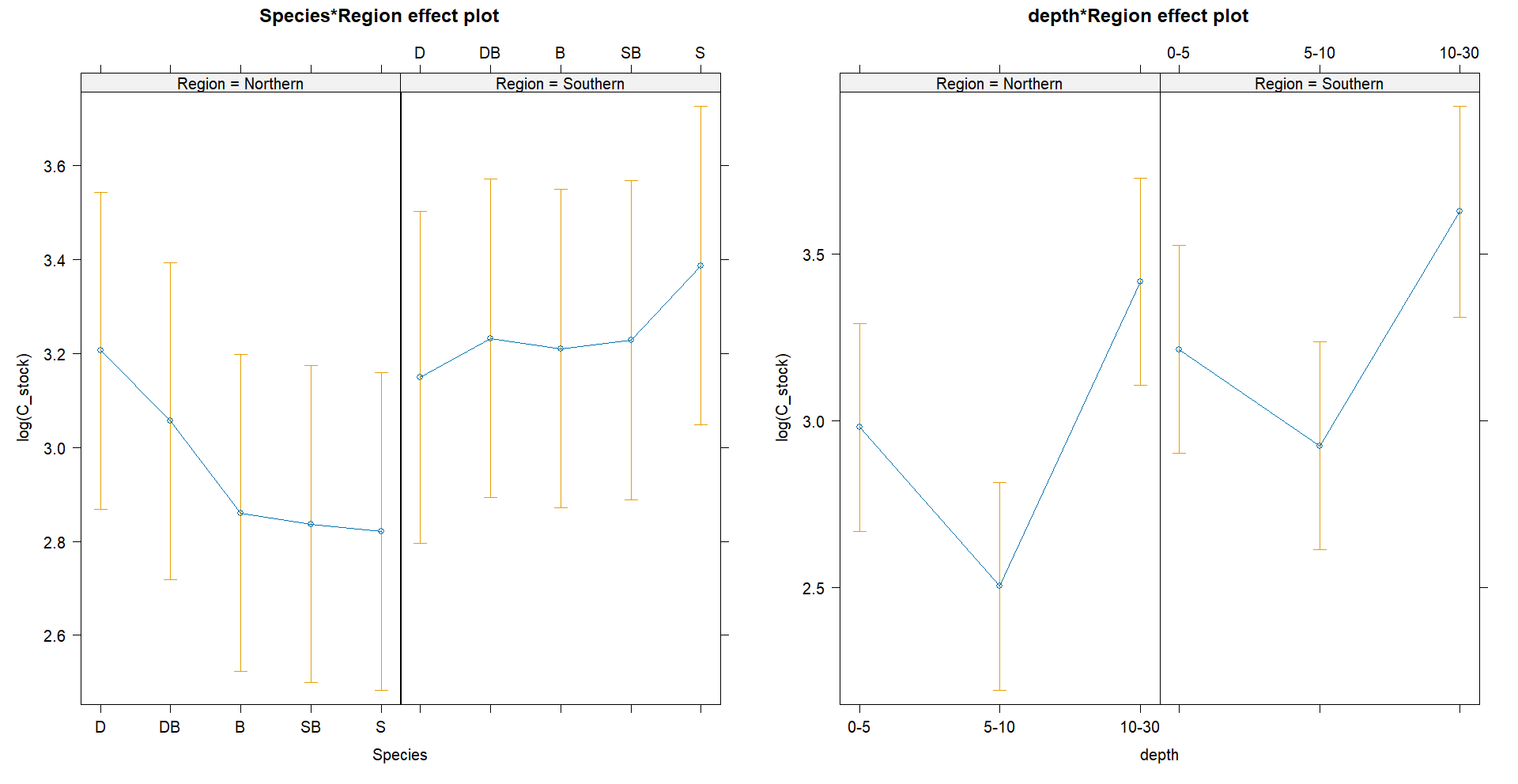

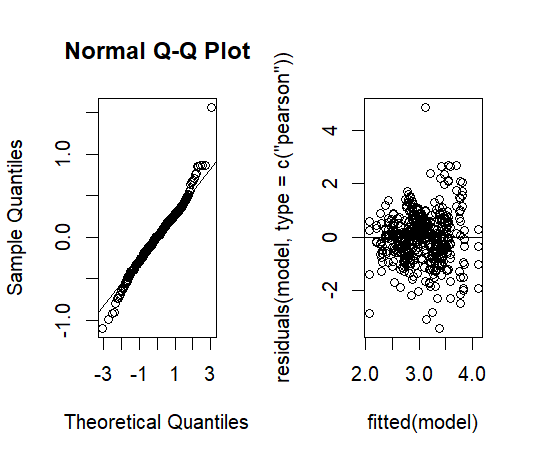

Figure S7. Linear mixed model fitted model for C stocks.

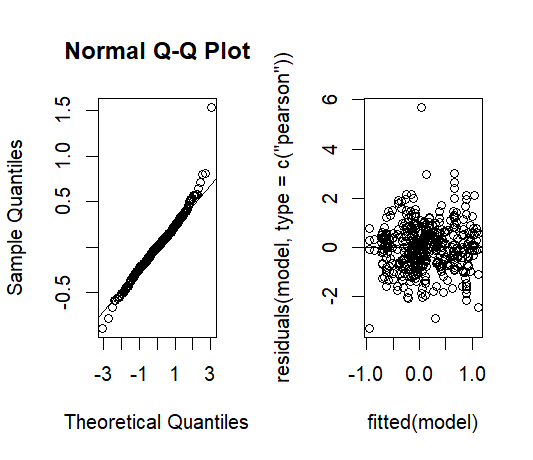

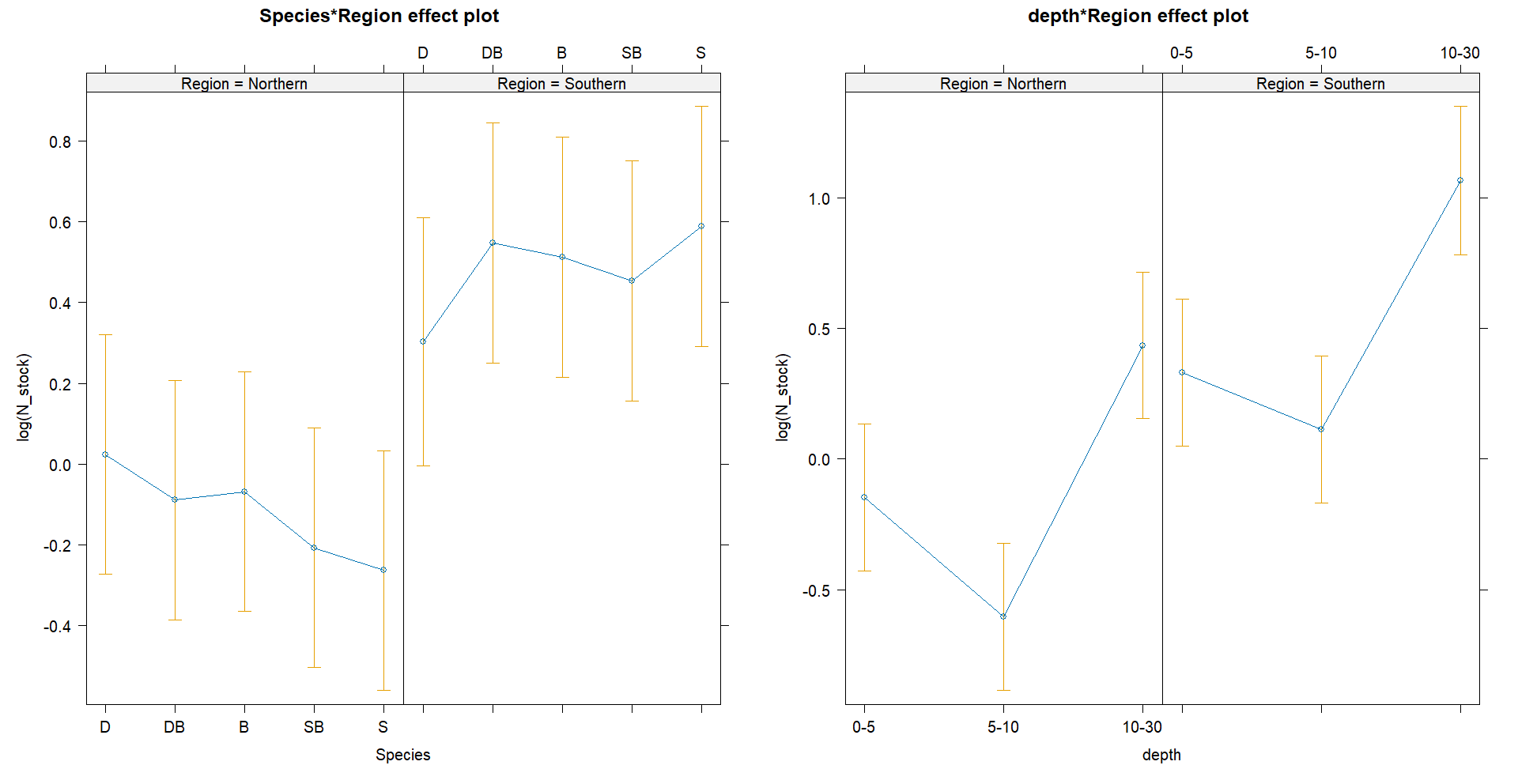

Figure S8. Linear mixed model fitted model for N stocks

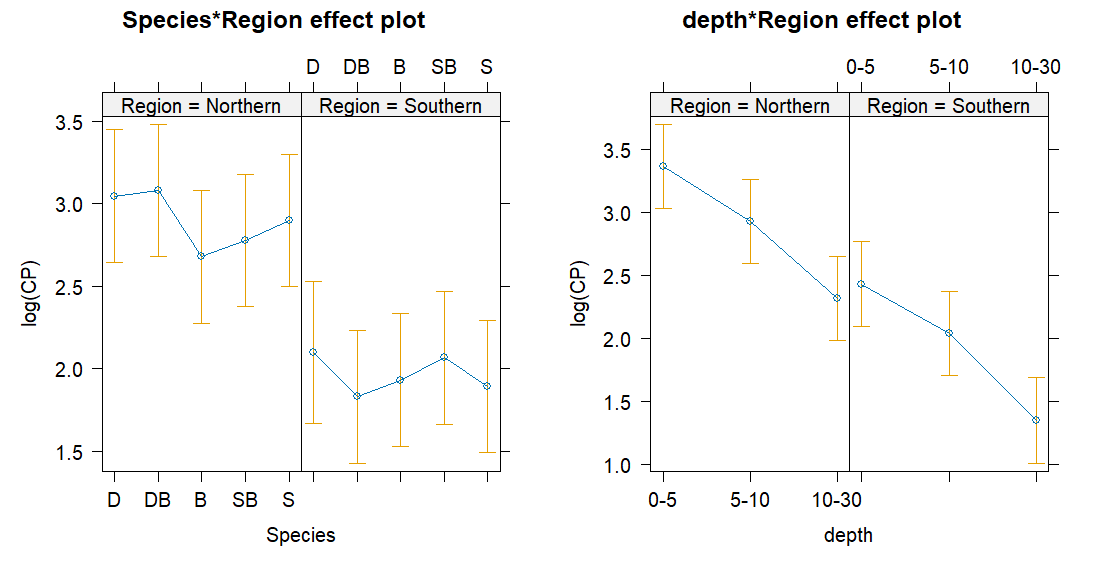

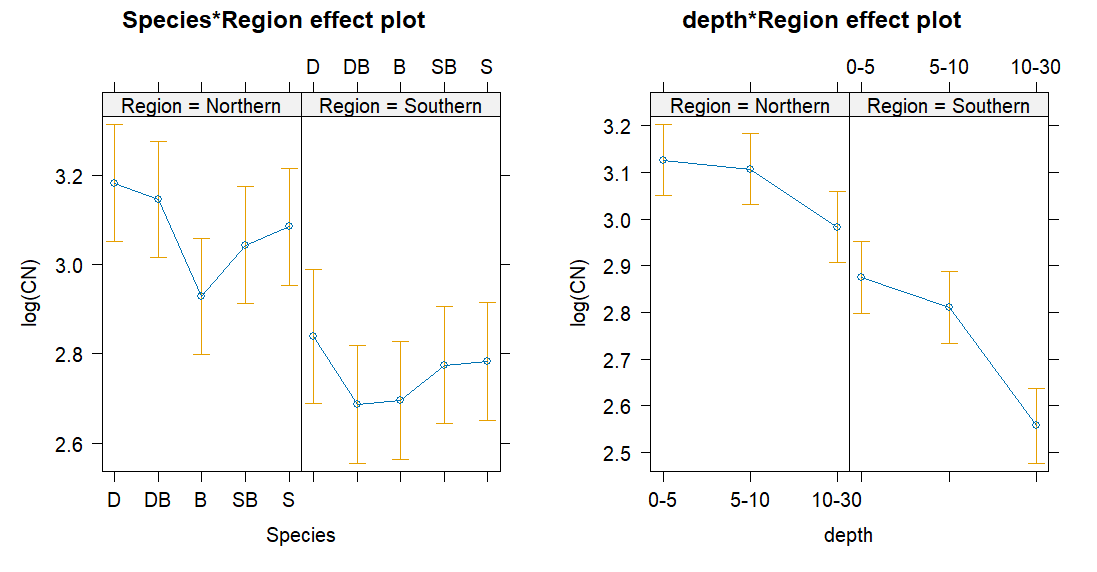

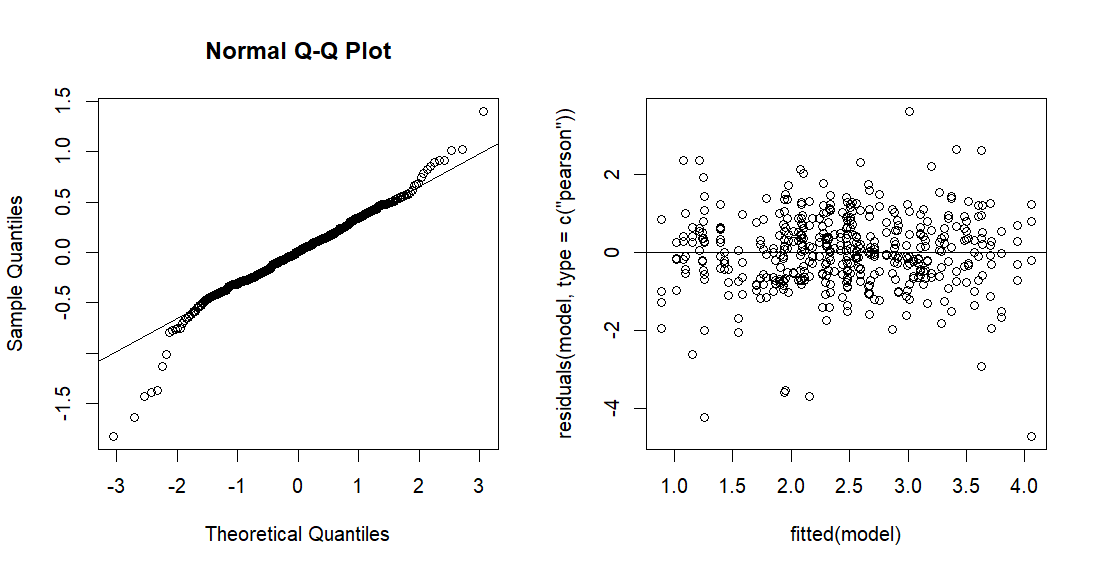

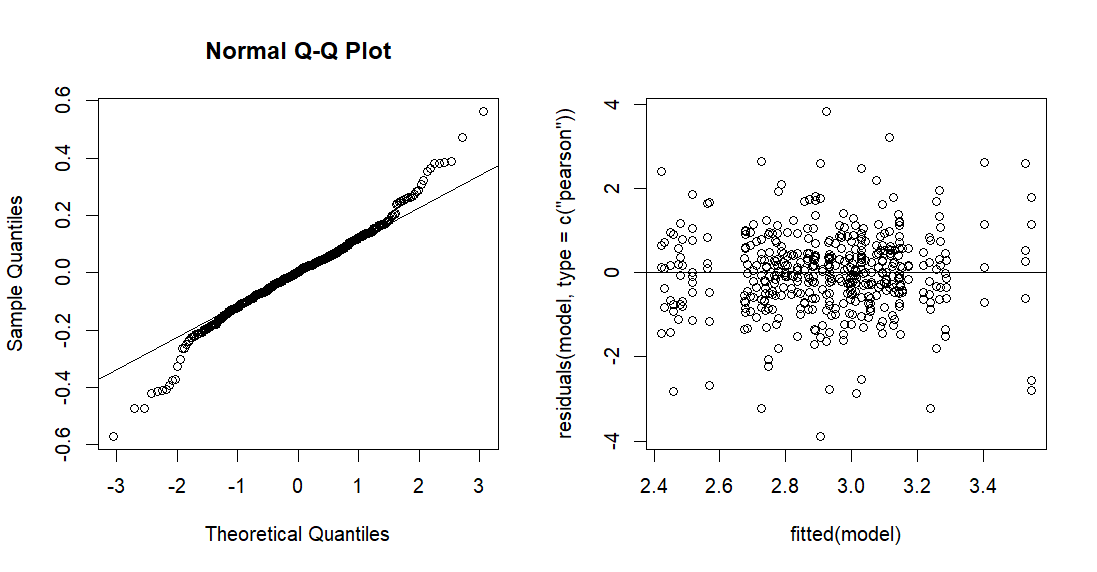

Figure S9. Linear mixed model fitted model for stochiometric ratios (CN and CP) for the organic layer.
